# Multimodal characterization of antigen-specific CD8^+^ T cells across SARS-CoV-2 vaccination and infection

**DOI:** 10.1101/2023.01.24.525203

**Authors:** Bingjie Zhang, Rabi Upadhyay, Yuhan Hao, Marie I. Samanovic, Ramin S. Herati, John Blair, Jordan Axelrad, Mark J. Mulligan, Dan R. Littman, Rahul Satija

**Author notes:** These authors contributed equally (Bingjie Zhang, Rabi Upadhyay).

## Abstract

The human immune response to SARS-CoV-2 antigen after infection or vaccination is defined by the durable production of antibodies and T cells. Population-based monitoring typically focuses on antibody titer, but there is a need for improved characterization and quantification of T cell responses. Here, we utilize multimodal sequencing technologies to perform a longitudinal analysis of circulating human leukocytes collected before and after BNT162b2 immunization. Our data reveal distinct subpopulations of CD8^+^ T cells which reliably appear 28 days after prime vaccination (7 days post boost). Using a suite of cross-modality integration tools, we define their transcriptome, accessible chromatin landscape, and immunophenotype, and identify unique biomarkers within each modality. By leveraging DNA-oligo-tagged peptide-MHC multimers and T cell receptor sequencing, we demonstrate that this vaccine-induced population is SARS-CoV-2 antigen-specific and capable of rapid clonal expansion. Moreover, we also identify these CD8^+^ populations in scRNA-seq datasets from COVID-19 patients and find that their relative frequency and differentiation outcomes are predictive of subsequent clinical outcomes. Our work contributes to our understanding of T cell immunity, and highlights the potential for integrative and multimodal analysis to characterize rare cell populations.

## INTRODUCTION

The COVID-19 pandemic has represented an unprecedented challenge for global public health, but mRNA vaccines have demonstrated strong clinical efficacy in protecting against severe disease^1–3^. Immune responses elicited by SARS-Cov-2 mRNA vaccines are typically assessed via titers of B cell-derived neutralizing antibodies, which rise rapidly after vaccination boosts but decline after 3-6 months^4–7^. However, it is evident that cellular immunity, mediated in part by CD4^+^ and CD8^+^ T cells, plays a critical role in viral clearance and protection^8^. In particular, vaccine-induced T cells have been found to provide protection against COVID-19 even in the absence of antibody responses^9^, and cellular immunity is likely to play an important role in providing protection against new viral variants^10^. A deeper understanding of the distinct subpopulations that drive cellular immunity, as well as their molecular programs and developmental determinants, will be essential for interpreting individual immune responses and for informing public health strategies^11^.

Antigen-specific T cells are conventionally identified by profiling cytokine secretion or expression of activation-induced surface markers after *ex vivo* antigen exposure, or, alternately, by labeling with peptide-MHC multimers. Both types of assays can be multiplexed with additional surface proteins for flow cytometry assays^12^. Multiple studies have applied these approaches to profile responses to SARS-CoV-2 mRNA vaccination, focusing in particular on the kinetics of antigen-specific T cell proliferation, alongside surface marker characterization^7,8,13–19^. Longitudinal profiling of human peripheral blood mononuclear cells (PBMC) followed by pMHCI-tetramer enrichment revealed a clear induction of antigen-specific CD8^+^ T cells after prime and boost vaccination, followed by a subsequent contraction phase as cells differentiated in the subsequent 3-4 months^8^. *Ex vivo* activation experiments demonstrated similar kinetics, and highlighted the potentially limited sensitivity of these assays to quantify rare CD8^+^ cells^5,7,20^.

Single-cell RNA-sequencing (scRNA-seq) assays are, in principle, well-suited for characterization of cellular responses. They can complement flow cytometry to provide transcriptome-wide molecular readouts, providing rich information for phenotyping high-resolution stages as well as activation and differentiation trajectories^21–23^. Moreover, single-cell sequencing assays enable unsupervised identification of cell states directly from PBMC samples, without need for *ex vivo* restimulation to reveal pre-established immunophenotypic markers of differentiation, or specificity for particular HLA haplotypes. However, even as scRNA-seq assays increase in sensitivity and throughput, it can be challenging to detect rare or transcriptomically subtle cell states from sparse and noisy datasets. For example, a previous study that utilized scRNA-seq to profile COVID mRNA vaccine responses showed substantial activation and proliferation within myeloid clusters, but did not specifically identify antigen-specific T cell subsets^24^.

Here, we perform a longitudinal analysis of human PBMC from a SARS-CoV-2 mRNA vaccination time series using a suite of multimodal single-cell sequencing technologies. Moving beyond the transcriptome, we additionally measure (either in parallel or in separate experiments) chromatin accessibility, surface protein abundance, immune receptor repertoires, and pMHC/multimer-binding modalities. By leveraging computational tools for within- and across-modality integration, we identify specific groups of vaccine-induced CD8^+^ effector memory T cells in each dataset. This strategy enables us to delineate high-resolution subpopulations and biomarkers within each modality, validate their clonal identity and antigen-specificity, and identify their developmental regulators. Moreover, by integrating our datasets with single-cell datasets of natural SARS-CoV-2 infection, we track the temporal differentiation patterns of these cells and show that their quantitative abundance is strongly associated with recovery from severe disease.

## RESULTS

### Multimodal identification of vaccine-responsive CD8^+^ T cell subsets

To investigate immune responses to SARS-CoV-2 mRNA vaccination at single-cell resolution, we recruited an initial set of six healthy donors and analyzed circulating PBMC samples over a time course of BNT162b2 mRNA vaccination. Specimens were collected in the first four months of 2021 from donors with no self-reported previous experience with SARS-CoV-2 infection (Supplementary Methods, Supplementary Table 1). Donors were profiled at four time points: immediately before (Day 0) vaccination, after primary vaccination (Day 2, Day 10), and seven days after boost vaccination (Day 28). For each of the 24 samples, we performed two multimodal single-cell sequencing assays: CITE-seq for simultaneous measurement of cellular transcriptomes and surface proteins^25^, and ASAP-seq for simultaneous profiling of open chromatin regions alongside cell surface proteins^26^ (Figure 1A). For each assay, we utilized an optimized panel of oligo-conjugated antibodies (‘TotalSeq-A’ panels from BioLegend, Supplementary Table 2) along with the inclusion of additional markers. Our initial dataset represented 192,574 single cells in total.

**Figure 1:**
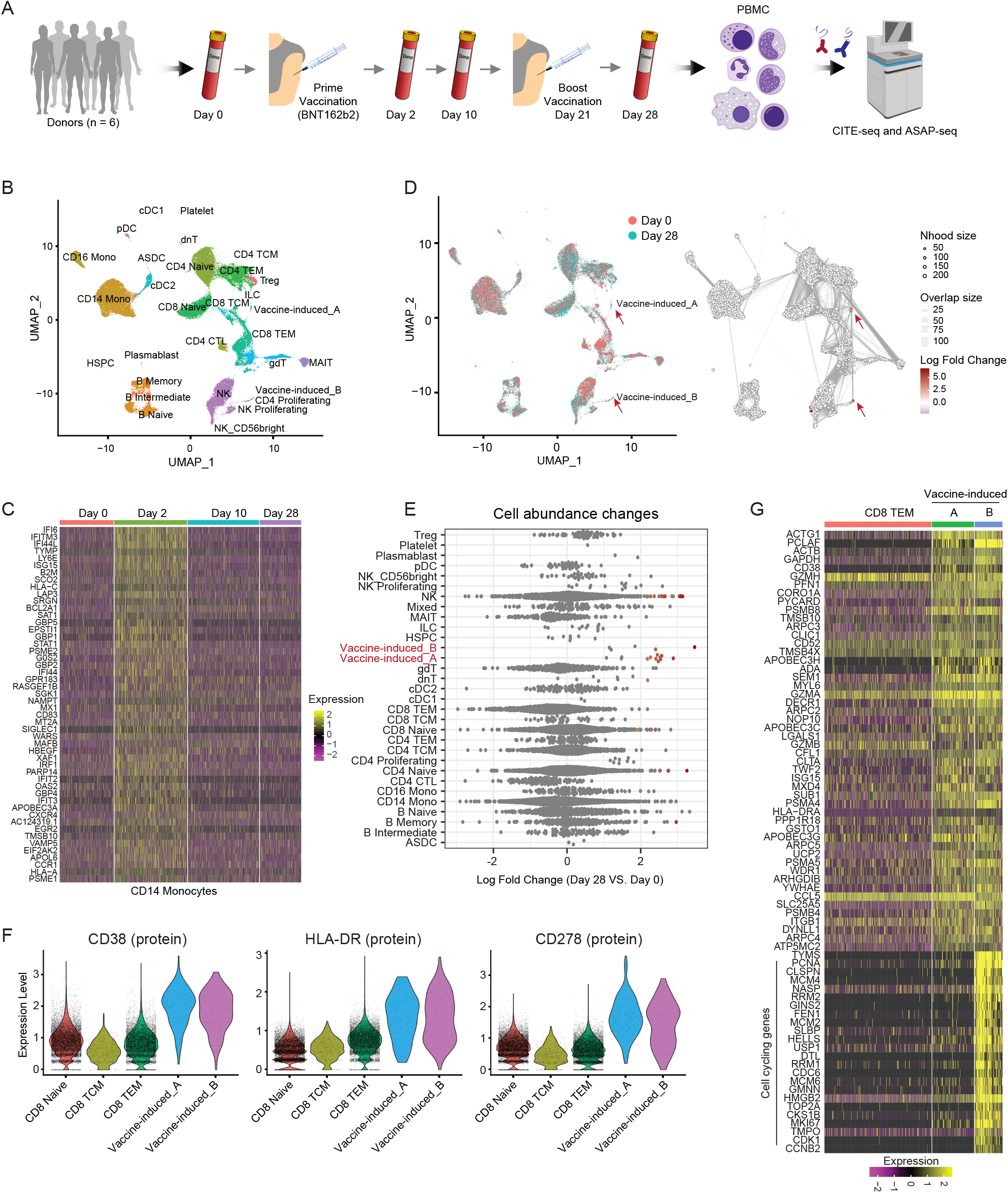
Multimodal identification of SARS-CoV-2 mRNA vaccine-induced CD8+ T cells. **(A)** Overview of human biospecimen study design. **(B)** Uniform manifold approximation and projection (UMAP) visualizations of 113,897 single cells profiled with CITE-seq and clustered by a weighted combination of RNA and protein modalities. Cells are colored based on level-2 annotation (level-1 and level-3 annotations are shown in Supplementary Figure 1A). **(C)** Single-cell heatmap showing activation of interferon response module within CD14 monocytes. **(D)** Milo analysis of differentially abundant cell states between Day 0 and Day 28 samples. UMAP on the left is color coded by time point. Right plot indicates embedding of the Milo differential abundance. Each node represents a neighborhood, node size is proportional to the number of cells, and neighborhoods are colored by the level of differential abundance. **(E)** Beeswarm plot showing the log-fold distribution of cell abundance changes. Neighborhoods overlapping the same cell population are grouped together, neighborhoods exhibiting differential abundance are colored in red. **(F)** Violin plots with protein upregulation of CD38, HLA-DR and CD278 (ICOS) in vaccine-induced cells compared with other selected CD8 T cells. **(G)** Heatmap showing mRNA expression of 50 marker genes for vaccine-induced group A cells, as well as cell cycling genes highly expressed in vaccine-induced group B cells. For visualization purposes, a randomly selected subset of CD8+ TEM are presented.

We first explored our CITE-seq datasets, applying our ‘anchor-based’ integration workflow to match together cells in shared biological states across individuals and time points^27,28^. Although this causes shared cell types in pre-vaccination and post-vaccination datasets to initially cluster together, integration enables us to consistently annotate these cell states across samples, and to learn cell type-specific responses in downstream analyses. To cluster cells, we applied ‘weighted-nearest neighbor’ (WNN) analysis (Supplementary Methods), which defines cell states jointly based on a weighted combination of RNA and protein modalities^28^. As we have previously shown^28^, WNN analysis improved the identification of cell states for multimodal technologies such as CITE-seq, by simultaneously leveraging the unsupervised nature of transcriptomic data with the robust protein measurements from oligo-tagged CITE-seq antibodies. We annotated clusters at three different levels of resolution (Figure 1B and Supplementary Figure 1A).

Comparing sample expression profiles across timepoints, we observed a strong activation of interferon-signaling pathways originating at the first post-vaccination time point (Day 2) and dampened by later time points, which was consistent with previous studies^5,24,29^ (Figure 1C and Supplementary Figure 1B). This response was most strongly activated in innate immune response components, but was weakly detectable in lymphoid cell types as well (Supplementary Figure 1C). The mRNA vaccine-responsive gene set was accompanied by the clear up-regulation of cell surface protein biomarkers including CD64 and CD169 in myeloid cell types^28^ (Supplementary Figure 1D).

We next explored changes in cellular density and abundance across our four vaccination timepoints. Strikingly, we identified two subsets of CD8^+^ T cells (‘vaccine-induced group A’ and ‘vaccine-induced group B’; Figure 1D-E) that were minimally present in Day 0 samples but increased moderately in abundance after primary vaccination, and sharply in abundance at Day 28 after boost vaccination) across multiple donors (Supplementary Figure 1E-F). We observed consistent results using either cluster-based differential abundance testing or alternately, using Milo, a framework for identifying differences in cellular density without reliance on cellular labels^30^. We observed only mild changes in cellular density among CD4^+^ T cell subgroups, likely due to earlier sampling timepoints in our experiment and the differential kinetics of CD4^+^ and CD8^+^ T cell responses^6,8^.

Both vaccine-responsive CD8^+^ T cell subsets exhibited up-regulation of protein biomarkers previously associated with activation during antigen-specific responses^8,31^, including CD38, HLA-DR, and CD278 (ICOS) (Figure 1F; additional markers in Supplementary Figure 1G). Including protein data using WNN analysis was essential for identifying and defining these subgroups, as they were not readily identifiable using unsupervised analysis of the transcriptomic data alone. Once identified, differential analysis revealed that group A and group B cells differed primarily in the expression of cell cycle genes (Figure 1G), while a module of 197 genes were consistently up-regulated across both groups (Figure 1G, Supplementary Figure 1H and Supplementary Table 3). This module was enriched for cytotoxic effector genes (GZMH, GZMA and GZMB), and also included multiple deaminase proteins (such as APOBEC3H, APOBEC3G, APOBEC3C and ADA) that can introduce mutations as part of the antiviral response^32,33^ (Figure 1G and Supplementary Figure 1I). We note that the identification of these two groups not only indicates the presence of both proliferative and non-proliferative populations, but also enables us to discriminate between proliferative responses (unique to group B), and activation responses (shared between groups A and B), which might otherwise blend together.

For additional validation, we re-analyzed a previously published CITE-seq dataset profiling a similar SARS-CoV-2 mRNA vaccination time course across six individuals^24^. While the original study did not identify populations of vaccine-induced CD8^+^ T cells in unsupervised transcriptomic analysis, we reasoned that supervised reference mapping workflows may have higher power to detect subtle cell states. Indeed, when mapping the query onto our newly generated reference, we identified both vaccine-induced populations (Supplementary Figure 2A). These cells sharply increased in frequency after boost vaccination (Supplementary Figure 2B), exhibited up-regulation of CD38 and ICOS surface protein levels, and were highly enriched in their expression of our identified vaccine-induced gene expression module (Supplementary Figure 2C). Taken together, we conclude that our multimodal analysis identifies CD8^+^ T cell subpopulations and molecular signatures that are induced after vaccination and are reproducible across donors and studies.

### Characterizing the epigenetic landscape of vaccine response

We next aimed to characterize vaccine-responsive programs defined by changes in chromatin accessibility. While transcriptomic measurements are rich descriptors of a cell’s current state and molecular output, ATAC-seq profiles are uniquely suited for identifying enhancers that exhibit heterogeneous activity, and for identifying regulators that establish and maintain cellular state. Our collected ATAC-seq profiles were obtained on the same biological samples as our CITE-seq data, but were collected from different cell aliquots. Particularly given the challenges in identifying and annotating high-resolution cellular states from scATAC-seq profiles^34,35^, we aimed to integrate chromatin accessibility profiles with our CITE-seq measurements.

To integrate datasets across modalities, we applied our recently developed ‘bridge integration’ approach, which performs integration of single-cell datasets measuring different modalities by leveraging a multi-omic dataset as a bridge^36^. We have previously demonstrated how this procedure can successfully map scATAC-seq query datasets onto scRNA-seq references using a publicly available “10x Multiome” dataset as a bridge^36^. Applying this workflow to our ASAP-seq datasets (Supplementary Methods), we annotated chromatin accessibility profiles by transferring labels from our CITE-seq reference (Figure 2A). We validated our inferred annotations using cell surface protein data that is simultaneously generated during ASAP-seq (Supplementary Figure 3A). For example, predicted monocytes were uniquely enriched for CD14 surface expression, predicted B cells exhibited CD19 upregulation, predicted dendritic cells exhibited up-regulation of FCER1A, and predicted CD8^+^ T and CD4^+^ T cells correctly expressed their canonical surface markers (Supplementary Figure 3A).

**Figure 2:**
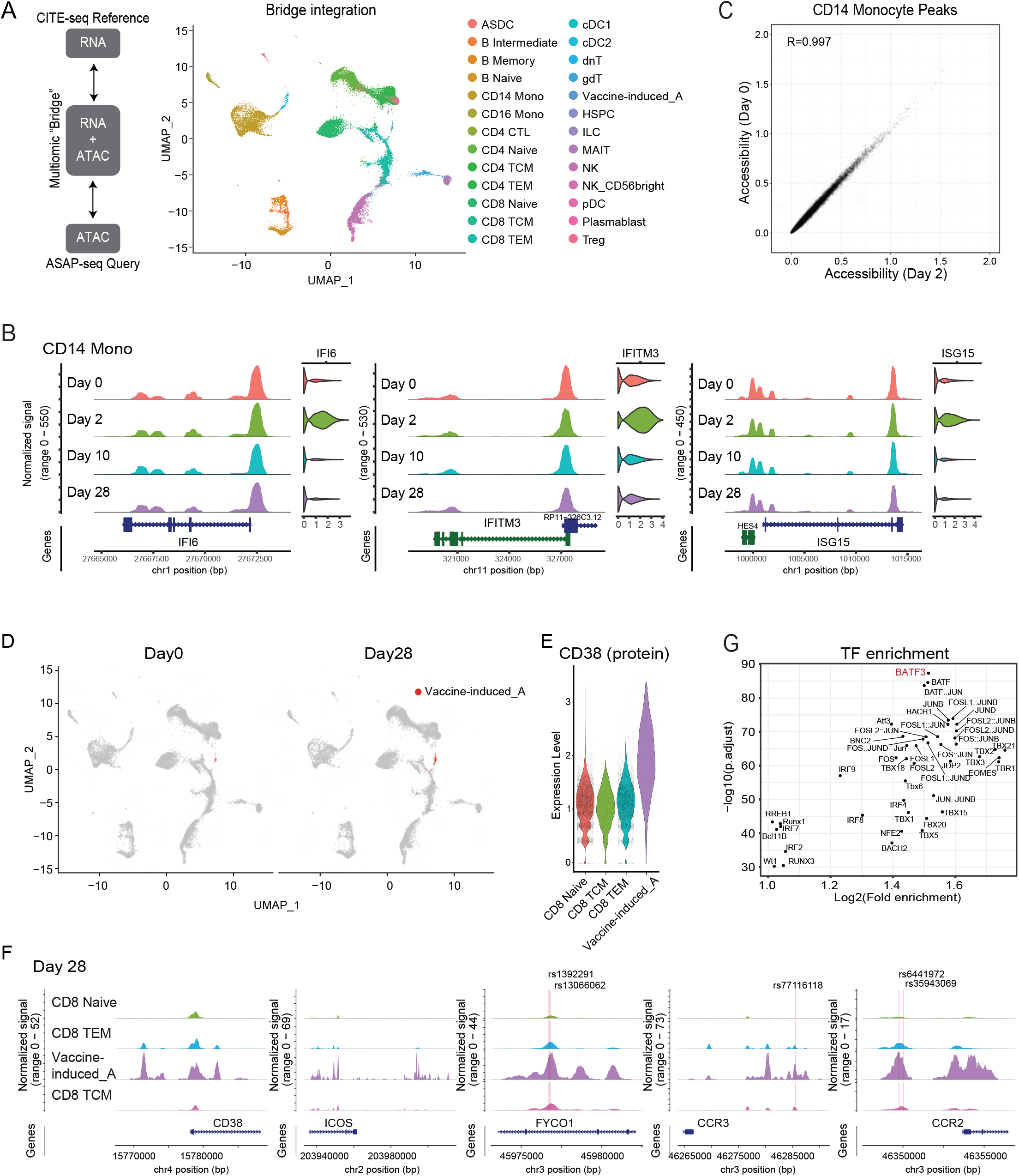
Celltype-specific chromatin accessibility dynamics in response to vaccination. **(A)** Bridge integration-based mapping of human PBMC scATAC-seq data onto the CITE-seq dataset from Figure 1B, using a multi-omic dataset as a bridge. Cells are colored by reference-derived annotation. **(B)** Coverage plots indicating chromatin accessibility around IFI6, IFITM3, and ISG15 in CD14 monocytes across all time points. Corresponding gene expression for each cell population, from the CITE-seq dataset, is shown on the right. **(C)** Scatter plot measuring correlation between Day 0 and Day 2 pseudo-bulk chromatin accessibility of CD14 monocytes. Each point corresponds to a called scATAC-seq peak. **(D)** UMAP visualization of scATAC-seq data on Day 0 and Day 28 after bridge integration. Vaccine-induced populations are highlighted in red. **(E)** CD38 protein expression levels, which were not considered during the bridge integration procedure, are correctly up-regulated in cells predicted to be vaccine-induced. **(F)** Examples of enhancer loci that are specifically accessible in vaccine-induced cells. Chromatin accessibility patterns on Day 28 are shown for four selected cell types. SNP sites are annotated as yellow lines. **(G)** Motif-based overrepresentation analysis of transcription factor binding sites in the top 1000 peaks with differentially enriched accessibility in the vaccine-induced group A cells.

We first examined accessibility changes in the innate immune response, where we had previously observed strong up-regulation of a module of interferon stimulated genes (ISG). Surprisingly, we did not observe dramatic remodeling of chromatin accessibility for myeloid cells before and after vaccination (Figure 2B-C and Supplementary Figure 3B-C). In a genome-wide analysis, which included both proximal and distal regions (Supplementary Methods), the chromatin accessibility profiles of CD14 monocytes were highly concordant before and after vaccination (Figure 2C, R=0.997). While we did detect a small number of peaks (n=75) that were differentially accessible across timepoints, these changes reflected minor quantitative fluctuations, as opposed to the opening or closing of regulatory regions (Figure 2C and Supplemental 3C).

These results suggest that the epigenetic landscape which is required to drive the transcriptional innate immune response is already established prior to vaccination, enabling the cells to quickly respond to external stimuli. We also identified nearly identical patterns when re-analyzing a published dataset^37^ of chromatin accessibility profiles before and after influenza vaccination (Supplementary Figure 3D-F; R=0.998). Taken together, these results demonstrate that chromatin accessibility patterns in myeloid cells exhibit only minor fluctuations during the initial innate immune response, and highlight how pre-established cell-type specific differences in accessibility correlate with future functional potential.

Notably, our bridge integration workflow also annotated vaccine-responsive CD8^+^ T cells in the ASAP-seq datasets. These cells increased sharply in frequency after boost vaccination (Figure 2D), and exhibited surface protein up-regulation of CD38, HLA-DR, and ICOS (Figure 2E and Supplementary Figure 4A). The surface protein measurements were not considered during the bridge integration procedure, and their consistency with the CITE-seq dataset represents an independent validation of our annotations. Moreover, these cells exhibited elevated gene ‘activities’ for the vaccine-induced gene module identified by CITE-seq (Supplementary Figure 4B). We did not observe a second population of proliferating cells in the ATAC-seq data, likely due to only subtle differences in chromatin accessibility that can accompany cell cycle changes^38^.

Having validated the identity of these cells, we next identified 2,678 peaks exhibiting differential accessibility (Supplementary Table 4), after comparison against other CD8^+^ T cell subsets (Supplementary Methods). These peaks included putative enhancer elements upstream of the CD38 and ICOS loci themselves, which are activated as cells acquire an activated state after immunization (Figure 2F). Globally, we found that 930 peaks were located near (within 20kb) genes that were up-regulated in vaccine-responsive CD8^+^ T cells. However, the remaining 1,170 peaks were located near genes that did not exhibit similar transcriptional differences, suggesting the pre-establishment of a chromatin landscape that will enable the downstream function of these cells. Interestingly, we found that enhancers specific to vaccine-specific cells harbored 13 SNPs that have been previously reported to be highly associated (p-value > 5e^-8^) with COVID susceptibility^39^, including within elements adjacent to FYCO1, CCR3, CCR2 and IFNAR2 (Figure 2F).

We next asked if the ASAP-seq data could reveal specific regulators required for the development and maintenance of vaccine-responsive subset. To accomplish this, we searched for transcription factor binding motifs that were overrepresented in specific peak subsets. We found that the motif for the transcriptional regulator BATF3 exhibited the strongest association with increased accessibility in vaccine-responsive CD8^+^ T cells (Figure 2G). While BATF3 has been characterized as a critical regulator of DC development^40,41^, recent studies in murine models have demonstrated its necessity for the specific development of CD8^+^ memory T cells^42^. Our findings extend these results to a human context, and suggest that our identified vaccine-induced subsets contribute to CD8^+^ T memory responses.

### Correlating molecular state with clonal identity

While our previous analyses identify and characterize CD8^+^ T cell populations that are induced in response to vaccination, our initial dataset cannot establish if these subgroups are mounting antigen-specific responses. To address this, we utilized dual DNA-oligo-tagged and fluorochrome-tagged peptide-class I MHC multimers^43^, constructed off a dextran backbone (“dextramers”, Figure 3A). We selected reagents designed to bind TCRs specific for immunodominant SARS-CoV-2 spike peptides, enabling direct *ex vivo* detection of antigen-specific T cells by either sequencing or cytometry. We selected 8 total donors carrying HLA-A*02:01 or HLA-B*07:02 alleles, and assayed for dextramer-positive (Dex^+^) cells initially by flow cytometry. We validated 5 such dextramer reagents to include in our panel (each targeting a separate peptide epitope), by demonstrating a robust and specific appearance of Dex^+^CD8^+^ T cells after vaccination (Supplementary Figure 5A).

**Figure 3:**
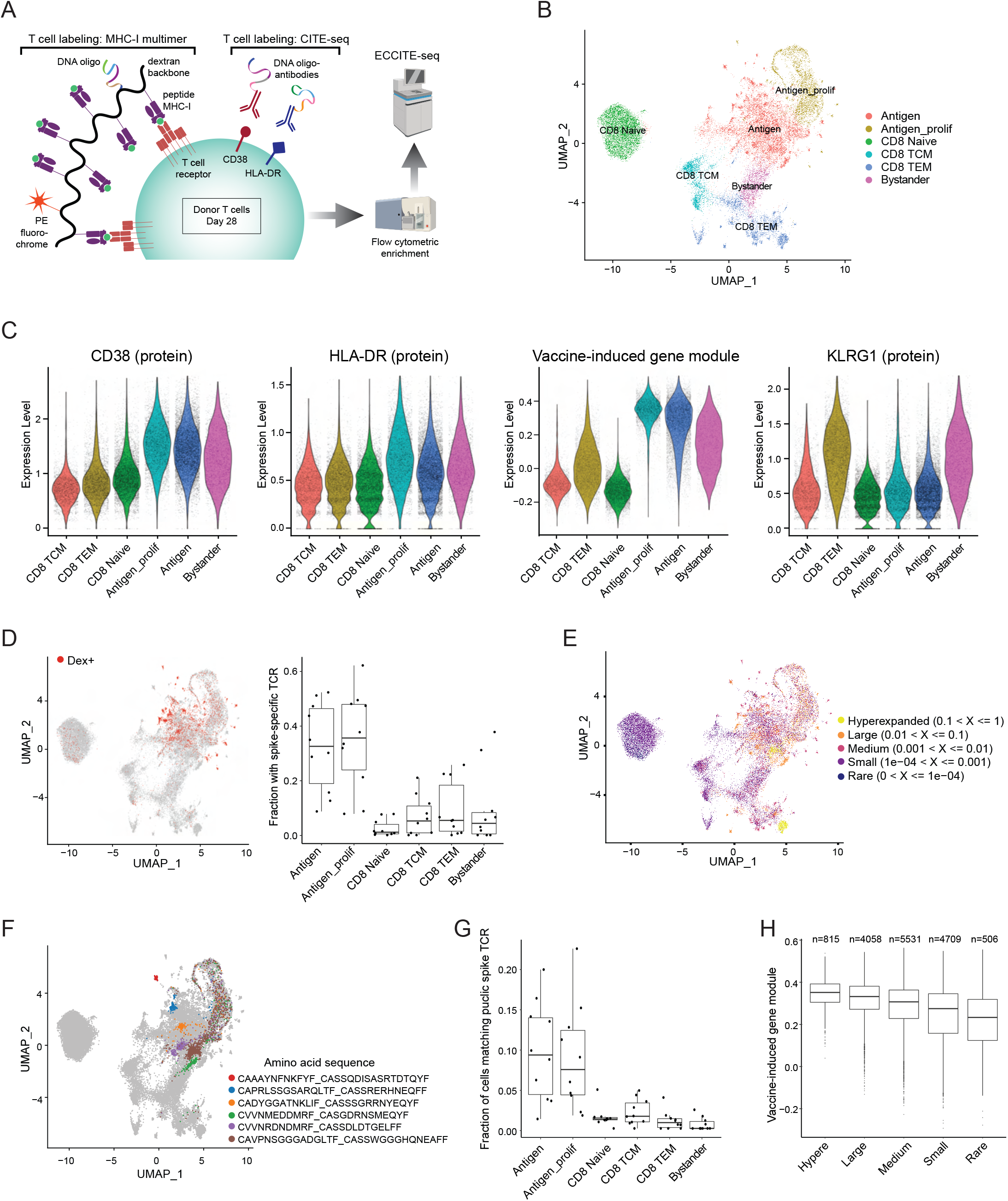
Antigen-specific clonal expansion of vaccine-induced CD8+ T cells. **(A)** Schematic of ECCITE-seq experimental design. **(B)** UMAP visualizations of 31,396 single cells profiled with ECCITE-seq and clustered based on weighted combination of RNA, protein and T-cell receptor information. **(C)** Violin plots for CD38, HLA-DR and KLRG1 protein levels, and the expression of identified vaccine-induced gene module. **(D)** Left: UMAP visualization from (B), dextramer-positive (Dex+) cells are highlighted in red. Right: The fraction of cells harboring spike-specific TCR in each cluster. A TCR clone is considered spike-specific when at least one cell of the clone is Dex+. Boxplot shows variation across n=10 samples. **(E)** UMAP visualization from (B), cells are colored by the expansion index of their associated clonotype based on TCR sequence information. **(F)** UMAP visualization from (B), cells representing the six most abundant spike-specific clones are highlighted.**(G)** Boxplots showing the fraction of cells harboring TCR matching SARS-CoV-2 spike antigens in public databases. **(H)** Boxplots showing the single-cell expression of the vaccine-induced gene module in antigen-specific cells. Cells are grouped by labels in (E).

To further explore heterogeneity within responding cells, we performed additional single-cell profiling using ECCITE-seq, which enables joint profiling of immunophenotypes, 5’-end transcriptomes, and immune repertoires^44^. We included the dextramer panel during antibody staining, enabling multiplexed detection of T cells specific for SARS-CoV-2 spike protein (Figure 3A). To enhance recovery of rare cell states, we restricted analysis to Day 28 PBMC cells and performed pre-enrichment steps via flow cytometric labeling and sorting, with 25% representing all CD8^+^ T cells, and 75% additionally enriched for CD38 expression and/or dextramer binding (Supplementary Methods). Our final dataset consisted of 31,396 single cells in total.

We clustered and visualized cells using WNN analysis based on three modalities (protein, transcriptome, T cell receptor sequence), allowing us to define cellular state based on all data types (Supplementary Methods). We identified six cell clusters, including CD8 naive cells and CD8 central memory subsets (Figure 3B). In addition, matching our CITE-seq dataset, we observed both cycling and non-cycling subsets of CD8^+^ T cells that exhibited elevated expression of our previously identified vaccine-induced gene module, as well as CD38 and HLA-DR surface proteins (“antigen” and “antigen_prolif”, Figure 3B-C). These clusters were strongly enriched for Dex^+^ cells (Figure 3D) as well as large and expanded cell clones (Figure 3E and Supplementary Figure 5B). We also found extensive TCR sharing between the cycling and non-cycling groups (Figure 3F). We therefore conclude that our identified vaccine-induced CD8^+^ T populations do represent antigen-specific cells that are responding to SARS-CoV-2 spike antigens.

Our enrichment strategy also enabled us to explore further sources of cellular heterogeneity amongst CD8^+^CD38^+^ T cells, which encompassed the clusters bearing our vaccine-induced gene signature. For example, we found that a subset of CD38^+^ cells uniquely expressed the inhibitory receptor KLRG1 surface protein (“bystander”, Figure 3B-C). In contrast to the two clusters populated by activated antigen-specific cells, CD38^+^KLRG1^+^ cells were not enriched for Dex^+^ cells (Figure 3D), they did not show evidence of expanded clonality, and they did not demonstrate enriched overlap with TCRs on antigen-specific cells (Figure 3E-F). To address the possibility that these CD38^+^KLRG1^+^ cells simply harbor TCRs not recognized by our dextramer panel, we also examined a large external database of TCRβ sequences^45,46^ specific for SARS-CoV-2 spike protein (Supplementary Methods). We found that unlike CD38^+^KLRG1^-^ cells, which demonstrated marked overlap with SARS-CoV-2 TCRs, the CD38^+^KLRG1^+^ population had minimal overlap with these documented clonotypes (Figure 3G, Supplementary Methods). These cells also exhibited weaker expression of the vaccine-induced gene module (Figure 3C), suggesting that CD38^+^KLRG1^+^ cells may represent cells harboring TCR with weak affinity for spike protein antigens, or alternatively, represent TCR-independent ‘bystander’ responses, such as those previously described within the microenvironments of tumors and other pathogens^47,48^.

Multiparameter flow cytometry on the Dex^+^ gate demonstrated that antigen-specific cells were indeed KLRG1^-^, in addition to being double positive for CD38 and HLA-DR, consistent with our initial CITE-seq (Supplementary Figure 5A). As these three markers represented prominent features from both our CITE-seq and ECCITE-seq experiments, we proceeded to gate for this population within all CD8^+^ T cells by flow cytometry and compare across time points (Supplemental Figure 5C-D). Validating our previous findings, we observed a striking induction of this population on Day 28—a result agnostic to the donor’s HLA haplotype or immunopeptidome (Supplemental Figure 5D). We conclude that CD8^+^CD38^+^HLA-DR^+^KLRG1^-^ cells are the most highly enriched for antigen-specific CD8^+^ T cells.

The rate of clonal expansion of antigen-specific T cells is an indicator of the strength of immune response^49^, and we therefore searched for gene expression patterns that were correlated with clonal size, even among antigen-specific cells. We found that the expression of the vaccine-induced gene module not only discriminated antigen-specific cells, but the module score also exhibited a dose-dependent relationship with clonal size (Figure 3H). We emphasize that the gene module is shared in both cycling and non-cycling groups, and therefore does not include proliferation-dependent genes that would be expected to correlate with clonal size. Instead, expression of this module likely reflects the signal strength of the original TCR-peptide interaction, an essential parameter which regulates the magnitude of clonal expansion and immune response^50–52^. Taken together, our multimodal ECCITE-seq dataset verifies the spike-specific nature of vaccine-induced CD8^+^ T cells, nominates specific biomarkers that subdivide heterogeneous activated populations, and identifies specific gene modules and surface markers which can be used to predict clonal dynamics, even in the absence of HLA haplotype and immune repertoire information.

### Identifying and tracing memory cells during COVID-19 progression

Having demonstrated that our identified gene modules could successfully infer the identity of antigen-specific cells across multiple vaccinated scRNA-seq datasets, we next asked if this signature was conserved in SARS-CoV-2 infected samples. We first examined a recent study that utilized a SARS-CoV-2 dextramer panel to identify long-lived memory cells during acute SARS-CoV-2 infection^53^. While unsupervised clustering of scRNA-seq data struggled to clearly identify dextramer-enriched cells (Figure 4A), we found that the expression of the vaccine-induced gene module had high predictive power (ROC = 0.88) to accurately predict dextramer staining labels (Figure 4B). Indeed, we found that our gene expression signature originally identified in vaccinated datasets was highly conserved in the dextramer positive cells (Figure 4C).

**Figure 4:**
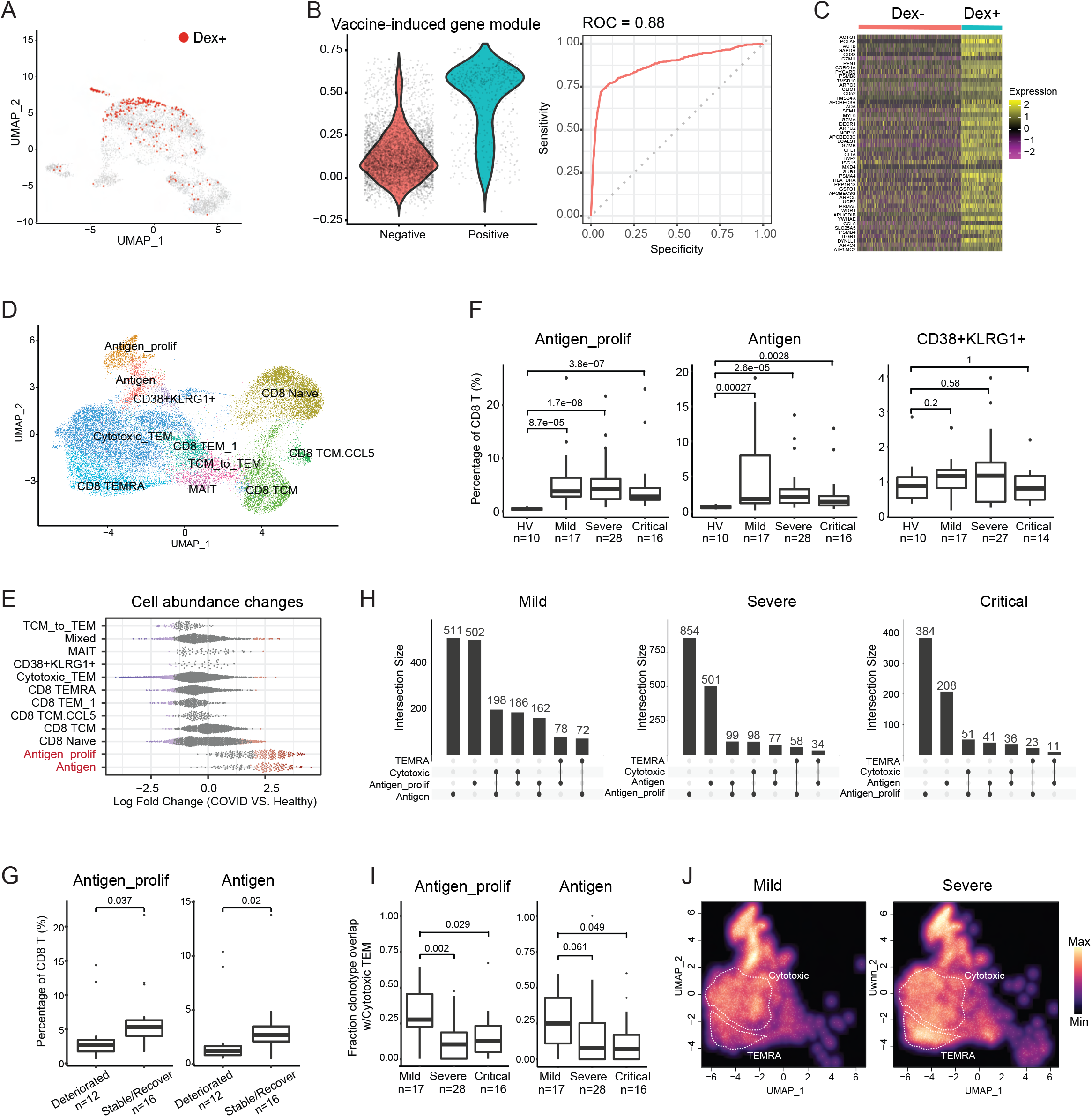
Inferred spike-specific T cells in SARS-CoV-2 infected samples. **(A)** UMAP visualization from (Adamo et al.), representing 6,070 CD8+ T cells collected during acute COVID-19 disease. Cells with positive dextramer staining are highlighted in red. **(B)** Left: Violin plots showing distribution of gene module score, comparing dextramer-positive versus -negative cells. Right: ROC curve assessing the ability of the gene module score to correctly predict dextramer staining labels in single cells. **(C)** The vaccine-induced gene expression module is conserved in SARS-CoV-2 infected samples. Shown is the expression of the top 50 vaccine-induced marker genes for dextramer-positive and -negative cells. For visualization purposes, a randomly selected subset of dextramer-negative cells are presented. **(D)** WNN UMAP visualization of 65,889 single cells from the COMBAT dataset. WNN was performed based on RNA and protein modalities, and identifies cell populations that we infer are specific to SARS-CoV-2 antigens. **(E)** Milo analysis of differential abundance changes, comparing healthy versus SARS-CoV-2 infected groups, as in Figure 1D-E. **(F)** Amongst all CD8+ T cells, donor fraction of antigen, antigen_proflif, and CD38+KLRG1+ cells, grouped by disease state. Boxplots show variation across n=71 donors, and p-values from two-tailed Wilcoxon signed-rank test. **(G)** The fraction of inferred antigen-specific cells correlates with clinical outcome. Same as in (F), but restricted to patients exhibiting severe symptoms, and grouped by their clinical outcome. **(H)** UpSet plot visualizing the overlap of antigen-specific TCR sequences across distinct molecular states of CD8+ T cells. **(I)** Fraction of TCR clonotypes identified in either antigen cells (right) or antigen_prolif cells (left) that are also observed in Cytotoxic TEM cells. Boxplots show variation across diseased donors. **(J)** Density plots showing the abundance distribution of all cells harboring expanded antigen-specific TCR sequences.

Multiple recent studies have reported that SARS-CoV-2-specific adaptive immune responses are associated with milder disease^54–56^. We therefore speculated that the abundance of CD8^+^ antigen-specific cells may correlate with disease phenotype and progression. To test this, we re-analyzed a large ECCITE-seq dataset (transcriptome + surface protein + TCR) from the COvid-19 Multi-omics Blood ATlas (COMBAT)^57^, which contains 65,889 CD8^+^ T cells prospectively collected from 10 healthy volunteers and 61 COVID-19 patients as they were admitted to inpatient hospital care, and who subsequently manifested mild, severe, or critical disease^57^. Applying our WNN integrative analysis pipeline, we identified analogous clusters enriched in the expression of our vaccine-induced gene module, as well as surface expression of CD38 and HLA-DR (Figure 4D and Supplementary Figure 6A). Based on our previous analyses (Figure 4B), we infer these populations to be specific to SARS-CoV-2 antigens. Abundances of both the cycling and non-cycling clusters (“antigen_prolif” and “antigen”) were sharply elevated in all infected samples, as compared to healthy controls (Figure 4E and Supplementary Figure 6B). Interestingly, we also identified subpopulations of CD38^+^ cells with heterogeneous surface expression of KLRG1 (Figure 4D and Supplementary Figure 6A).

We then examined the abundance of these identified populations and found that the CD38^+^KLRG1^-^ (both “antigen” and “antigen_prolif” clusters) but not the CD38^+^KLRG1^+^ clusters were associated with the severity and trajectory of COVID-19. We found that the relative abundance of predicted antigen-specific cells was sharply increased in diseased samples compared to healthy controls, but exhibited a progressive decrease across the spectrum of mild to critical severity patients (Figure 4F). We further explored the eventual outcome of 28 donors whose samples were collected during severe disease, and considered subsets of 16 patients who stabilized or recovered, versus 12 who further deteriorated. We found that the relative abundance of inferred antigen-specific cells was distinct between the two groups (Figure 4G). These data demonstrate that a mild clinical outcome is associated with an increased frequency of CD8^+^CD38^+^KLRG1^-^ T cells at the outset of illness, and suggest that patients who do not mount effective cellular immune responses are more likely to succumb to critical COVID-19.

We next explored relationships between immune repertoire sequences and molecular state, both of which were simultaneously measured in the COMBAT dataset. As expected, predicted antigen-specific cells were highly enriched for cells participating in either large- or hyper-expanded clones (Supplementary Figure 6C). We found that only CD8^+^CD38^+^KLRG1^-^ cells exhibited enriched overlap with a public database of SARS-CoV-2 TCR sequences (Supplementary Figure 6D), demonstrating that in both vaccination and infection, KLRG1 expression demarcates heterogeneous immune responses amongst activated and responding CD8 T cells.

Lastly, we observed extensive TCR sharing between different CD8^+^ T cell subsets, indicating evidence for lineage-specific differentiation trajectories. As we observed during vaccination, there was extensive clonal overlap between cycling and non-cycling antigen-specific cells (CD38^+^KLRG1^-^) (Figure 4H). Exploring overlap with differentiated cells, we found the most significant overlap with highly cytotoxic TEM (CD8^+^CD127^-^CD45RA^-^CD27^-^) subsets, with lower overlap with additional subsets including TEMRA cells (CD8^+^CD127^-^ CD45RA^+^CD27^-^), and effector cells with reduced cytotoxicity (CD8^+^CD127^mid^CD45RA^-^ CD27^mid^). Interestingly, we also found that the molecular state of differentiated T cells sharing CD38^+^KLRG1^-^ TCRs also varied as a function of disease severity. In samples with mild SARS-CoV-2 infection, we observed that nearly 25% of TCR sequences observed in predicted antigen-specific subsets exhibited clonal overlap with cytotoxic subsets of TEM (CD8^+^CD127^-^CD45RA^-^ CD27^-^), but this percentage was sharply reduced (in severe group: median of 7.74% for antigen cells, median of 10% for antigen_prolif cells; in critical group: median of 7.14% for antigen cells, median of 12.1% for antigen_prolif) in donors with severe and critical disease (Figure 4I). We did not observe similar findings for TEMRA cells (CD8^+^CD127^-^CD45RA^+^CD27^-^; Supplementary Figure 6E), and as a result, the distribution of cells harboring expanded antigen-specific TCR sequences was skewed towards TEMRA fates in these samples (Figure 4J and Supplementary Figure 6F). We confirmed that these findings were not driven by any potential correlation between disease severity and time since onset (Supplementary Figure 6G). Together, these results exhibit how the abundance and molecular differentiation outcomes CD38^+^HLA-DR^+^KLRG1^-^ during SARS-CoV-2 infection are predictive of disease severity and clinical progression.

## Discussion

Here, we present an extensive multimodal analysis of the immune response to SARS-CoV-2 mRNA vaccination using a suite of single-cell technologies. Our results emphasize the importance of multimodal technologies and datasets, particularly when characterizing rare populations. We found that a weighted combination of both protein and transcriptome features was essential for initial identification of antigen-specific CD8^+^ T cells, which had previously been unidentified in scRNA-seq datasets of vaccinated human donors. Moreover, layering additional molecular modalities onto our initial map deepened our understanding of these cells. Leveraging ‘bridge integration’ to annotate these cells in scATAC-seq datasets led to the discovery of numerous cell type-specific enhancers and transcriptional regulators that establish and maintain cellular state. Similarly, additional datasets that measured immune repertoires and the binding of MHC-I dextramers established the antigen-specificity and clonal dynamics of these populations. Even though no single technology allows simultaneous profiling of all molecular modalities, our integrated experimental design enabled us to deeply characterize these cells.

While our identified protein biomarkers CD38 and HLA-DR have been previously used to characterize antigen-specific CD8^+^ T cells in flow cytometry assays^53,58^, our unsupervised single-cell profiling strategy identified additional heterogeneity within this important subset. In addition to identifying both cycling and non-cycling subsets, we observed heterogeneity in the expression of KLRG1 within this group, and found that KLRG1^-^ subpopulations were most likely to contain highly clonal cells that exhibited binding to spike-specific dextramer reagents. While KLRG1 is a highly cytotoxic molecule, previous studies have linked its expression within antigen-specific memory cells to a short-lived phenotype^59–61^. Our results suggest that this surface marker distinguishes cells with distinct antigen specificities, which likely contributes to downstream differences in their phenotype and persistence.

By leveraging molecular signatures we identified within vaccinated samples, we were able to annotate antigen-specific CD8^+^ T cells in additional published datasets^24^, including those prospectively collected from COVID-19 patients^53,57^. In these samples, we also leveraged immune repertoire information to link antigen-specific CD8^+^ T memory precursors with their differentiated progeny. We found that disease severity and outcome correlated not only with the abundance of precursor cells, but also with the molecular state of their descendants, and in particular we found that donors who manifested extensive TCR sharing between memory precursors and cytotoxic progeny were associated with a milder clinical course. These results exemplify a potential mechanism by which cellular immunity may play an important role in resolving viral infection.

While our study is rooted in analyzing mRNA vaccination and coronavirus disease, the antigen-specific CD8^+^ T cell subpopulations we uncover are likely to represent features of human immune responses more broadly. For example, a recent study of neoadjuvant head and neck cancer immunotherapy patients identified a subpopulation of circulating CD8^+^ T cells, similarly enriched for CD38 and HLA-DR expression, whose abundance within the primary tumor and within PBMC changed after a 3-week course of checkpoint blockade therapy^62^. In a separate context, the study also identified heterogeneity in KLRG1 expression and found that the specific abundance of PD1^+^KLRG1^-^ cells within that subset positively correlated with optimal induction of tumor antigen-specific T cells and overall treatment outcome. Taken together, these results demonstrate the potential for monitoring of antigen-specific T cells to inform our understanding of disease and treatment trajectories.

## Data availability

Seurat and Signac are freely available as open-source software packages:

CITE-seq, ECCITE-seq, and ASAP-seq datasets generated for this manuscript are available at: https://zenodo.org/record/7555405

## Acknowledgements

The authors would like to thank all the members of the Satija and Littman labs for thoughtful discussions related to this work. B.Z. is a postdoctoral fellow of the Jane Coffin Childs Memorial Fund for Medical Research. This investigation has been aided by a grant from the Jane Coffin Childs Memorial Fund for Medical Research. R.U. is a Damon Runyon Physician-Scientist supported (in part) by the Damon Runyon Cancer Research Foundation (PST-25-19). This work was supported by the Chan Zuckerberg Initiative (EOSS5-0000000381, HCA-A-1704-01895 to R.S. and D.R.L.), the Howard Hughes Medical Institute (D.R.L.), and the NIH (AI082630, AI158617 to R.S.H; RM1HG011014-02, 1OT2OD033760-01 to R.S).

---

**Supplementary Figure 1.**
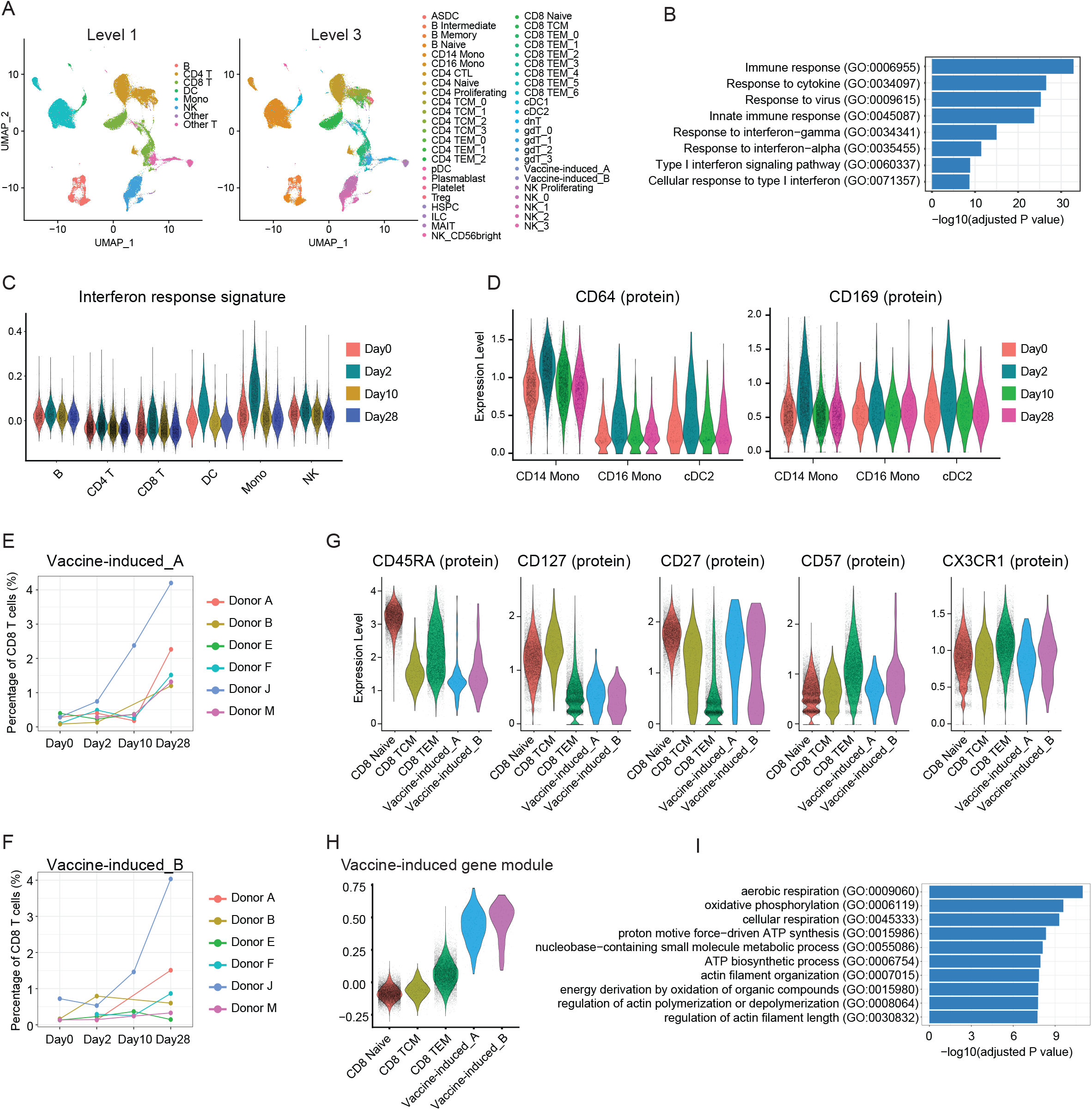
**(A)** UMAP visualizations of 113,897 single cells profiled with CITE-seq and clustered on the weighted combination of both RNA and protein modalities. Cells are colored with either level 1 or level 3 annotations. **(B)** Enriched GO terms for activated (Day 2 vs Day 0) genes in CD14+ Monocytes. **(C)** Violin plots of interferon response signatures in selected cell types across four timepoints. **(D)** Violin plots of protein upregulation of CD64 and CD169 in single cells in selected cell types, across four timepoints. **(E-F)** Percentage of CD8+ T cells in vaccine-induced groups for each donor across four timepoints. **(G)** Violin plots showing the protein expression of CD45RA, CD127, CD27, CD57 and CXC3R1 in selected cell types. **(H)** Violin plots comparing upregulated gene module scores in vaccine-induced group A and B cells, as well as selected other subsets. **(I)** Enriched GO terms for the 197 signature vaccine-induced gene set.

**Supplementary Figure 2.**
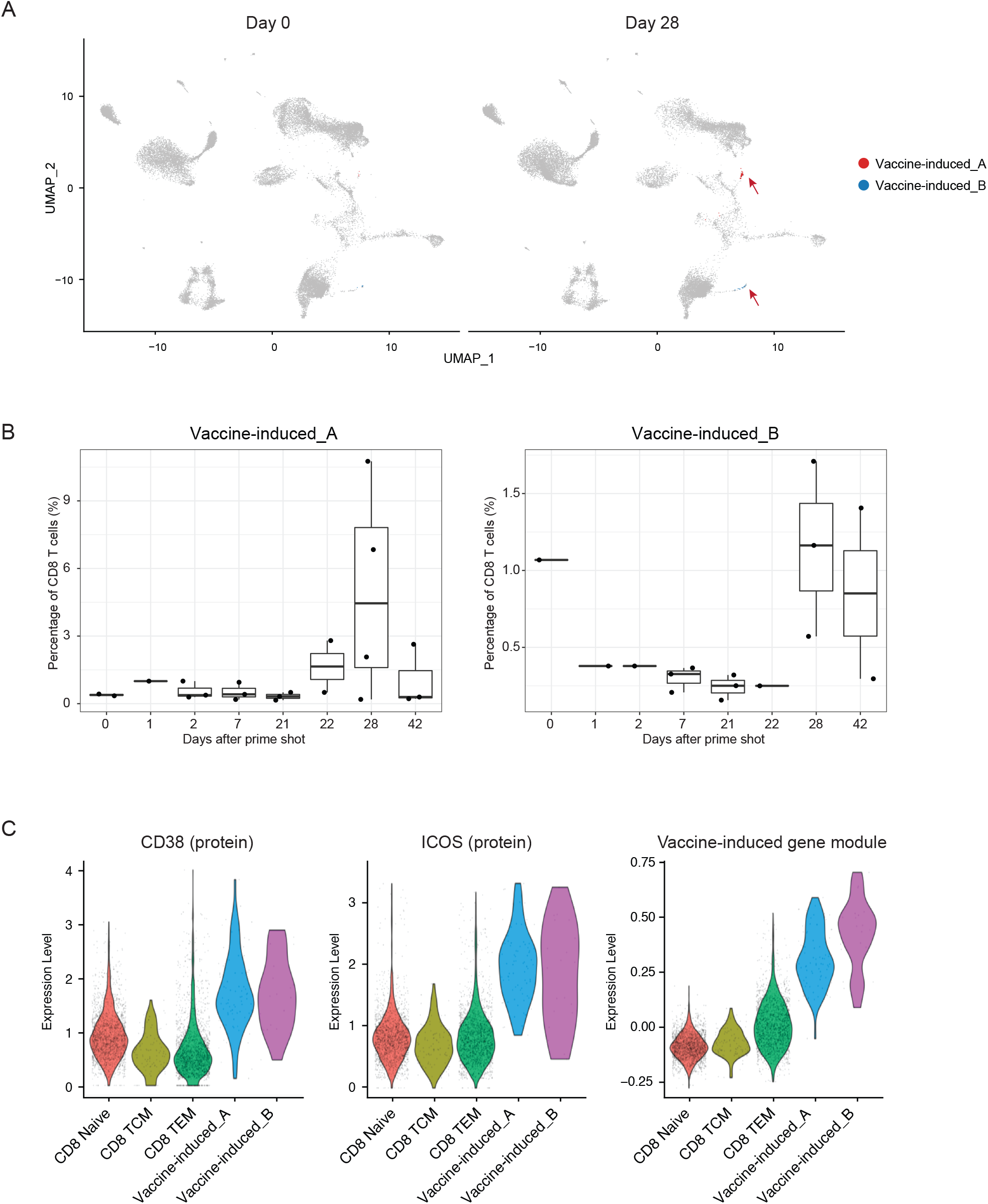
**(A)** UMAP visualization of CITE-seq data derived from human PBMC from Arunachalam et al. on Day0 and Day28, after reference mapping to the CITE-seq data in Figure 1B. Cells matching gene signature for vaccine-induced group A and B cells are highlighted in red and blue. **(B)** Boxplots showing the percentage of CD8+ T cells that fall in vaccine-induced group A (left) or group B cells (right) for each donor across eight timepoints. Each dot represents one donor. **(C)** Violin plots showing protein expression of CD38 and ICOS, along with a gene module score for vaccine-induced cells in selected CD8+ T cell subsets.

**Supplementary Figure 3.**
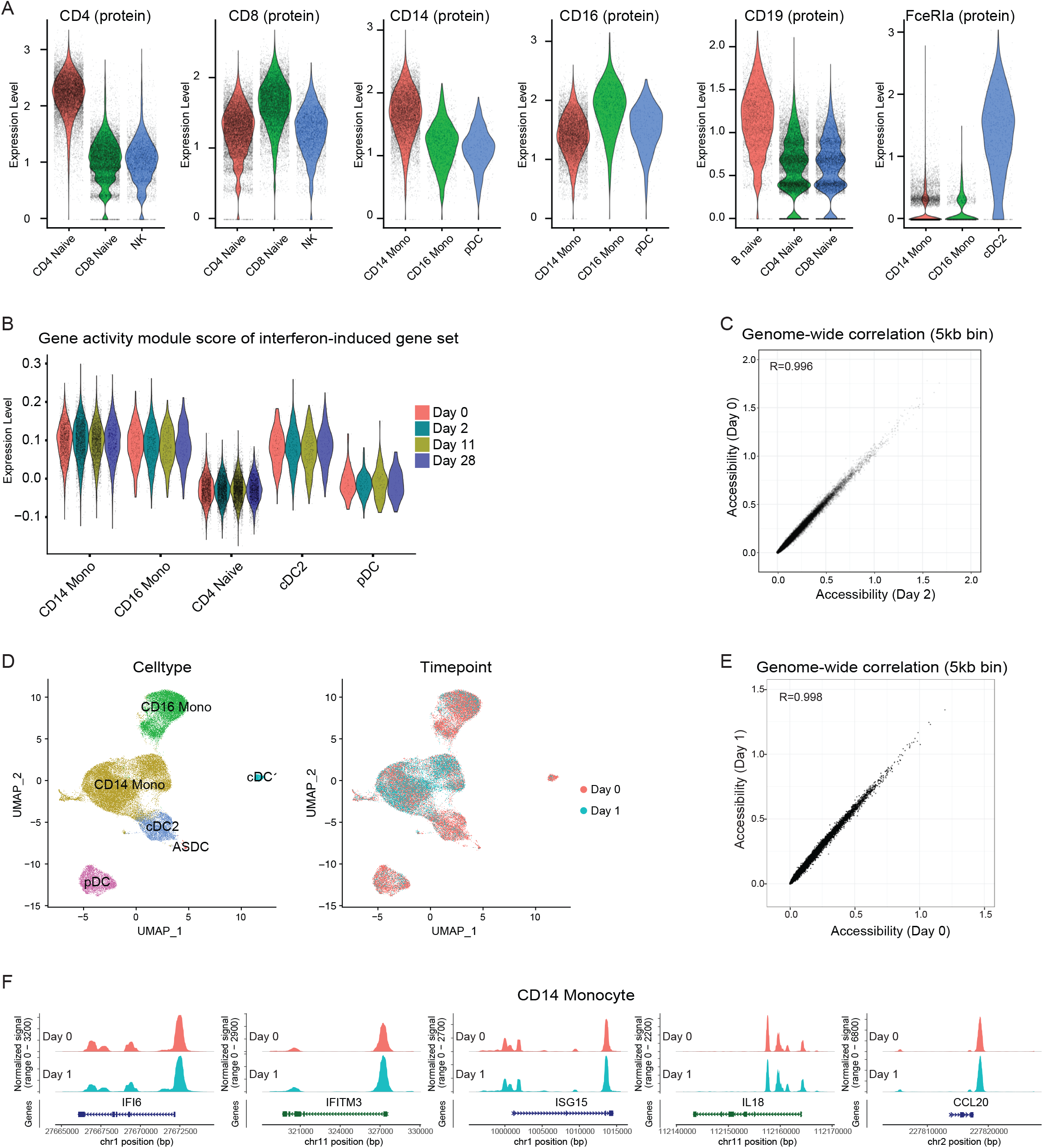
**(A)** Violin plots showing the expression of canonical surface proteins in the ASAP-seq dataset. Cells are grouped by bridge integration-derived labels. Proteins visualized include markers of CD4 and CD8 T cells, CD14 and CD16 monocytes, B cells and cDC2 cells. **(B)** Violin plots showing the gene ‘activity scores’, which are derived from scATAC-seq data, of the interferon-induced gene set shown in Figure 1C. **(C)** Scatter plot showing the correlation between pseudobulk chromatin accessibility of CD14 monocytes from Day 0 and Day 2 samples. Each point corresponds to a 5KB genomic bin. **(D)** UMAP visualization of the scATAC-seq profiles of myeloid cells from a trivalent inactivated seasonal influenza vaccine (TIV) study. Cells are colored by annotations (Left) or timepoints (right). **(E)** Scatter plot showing the correlation comparing pseudobulk chromatin accessibility of CD14 monocytes in the TIV dataset between Day 0 and Day 1. Each point corresponds to a 5KB genomic bin. **(F)** Visualization of open chromatin accessibility at representative loci on Day 0 and Day 1 in CD14 monocytes from the TIV study. Despite extensive transcriptional activation at these genes on Day 1, chromatin accessibility patterns do not change.

**Supplementary Figure 4.**
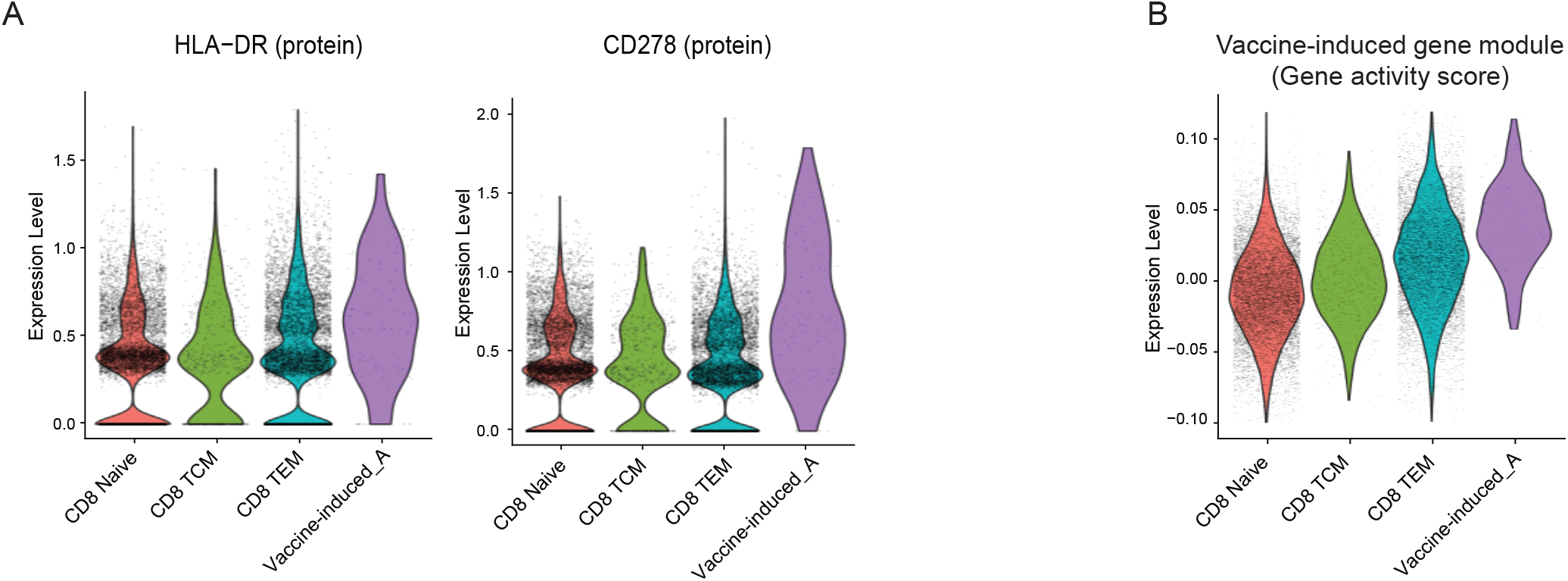
**(A)** Violin plots showing the protein upregulation of HLA-DR and CD278 (ICOS) in the vaccine-induced group A cells identified in the ASAP-seq dataset. Cells are grouped by their bridge integration-derived labels. **(B)** Violin plots showing the module score of the 197 signature vaccine-induced gene set in the ASAP-seq dataset. The module score is calculated based on gene activity scores, which are derived from scATAC-seq data.

**Supplementary Figure 5.**
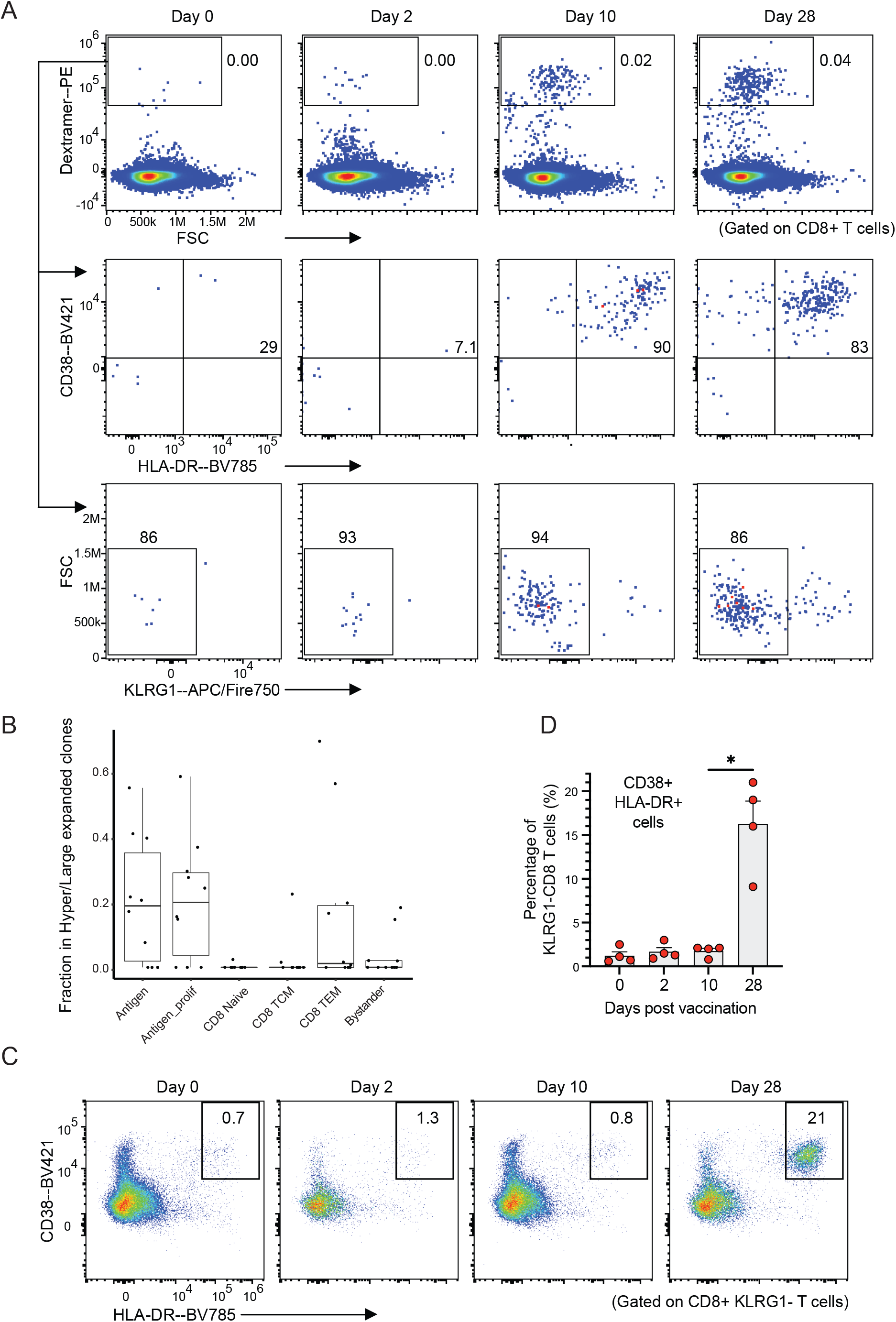
**(A)** Flow cytometry data generated during validation of individual dextramer reagents, with the progressive emergence of cells in the Dex+ gate across time points for a single donor. CD8+ cells were used as input. Middle and bottom row show CD38, HLA-DR, and KLRG1 abundance from the parent gate of Dex+CD8+ cells. **(B)** Boxplots indicate the fraction of cells harboring a hyper- or large-expanded TCR clone in each cluster. Each dot represents one biological sample. **(C)** Exemplary flow cytometry plots indicating the percentage of cells in CD38+HLA-DR+ gate of a single donor, from a parent gate of CD8+ KLRG1-cells. **(D)** Boxplot shows the percentage of CD38+HLA-DR+ cells in each donor, as a fraction of the CD8+KLRG1-gate exemplified in (C). Data represents n=4 donors with variable HLA haplotypes. p-value (p=0.0286) was calculated using a Mann-Whitney test.

**Supplementary Figure 6.**
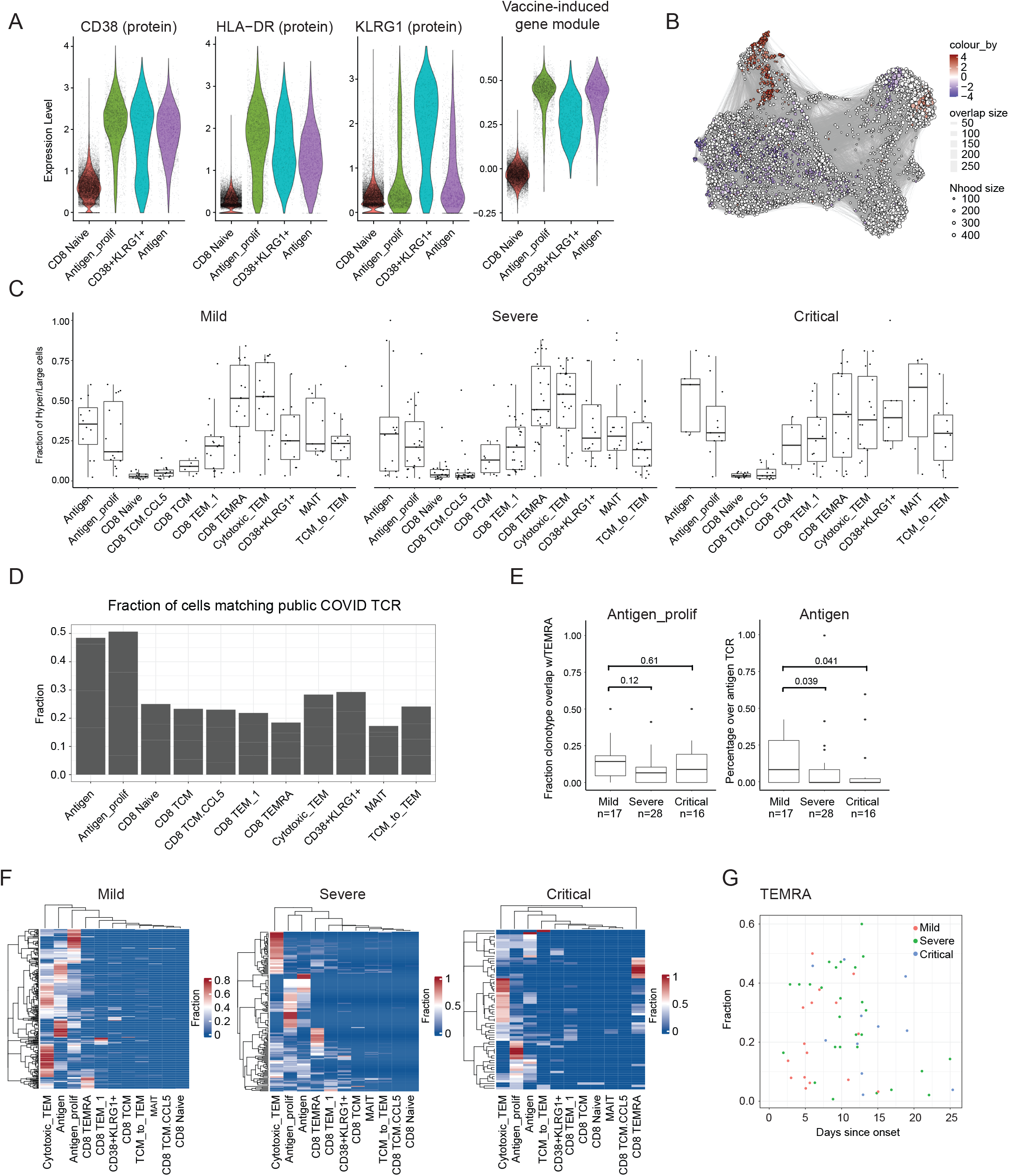
**(A)** Violin plots showing the protein expression of CD38 and HLA-DR, along with the signature gene module score for the vaccine-induced cells, in the COMBAT dataset. **(B)** Milo analysis of differential abundance changes between healthy and SARS-CoV-2 infected CD8+ T cells groups from the COMBAT dataset. UMAP visualization of the Milo differential abundance testing results. Each node represents a neighborhood. The size of nodes is proportional to the number of cells in the neighborhood. Neighborhoods are colored by their log fold changes for SARS-CoV-2 infected versus healthy groups. Only neighborhoods showing significant enrichment (SpatialFDR < 0.1 and logFC > 2) are colored. **(C)** Boxplots showing the fraction of cells harboring a hyper or large expanded TCR clone within each cluster. Each dot represents one biological sample. **(D)** Barplot showing the fraction of cells within each cluster harboring TCR matching SARS-CoV-2 antigens in public databases. **(E)** Fraction of TCR clonotypes identified in either antigen cells (right) or antigen_prolif cells (left), that are also identified in TEMRA cells. Boxplots show variation across diseased donors. **(F)** Heatmaps show the distribution of cells harboring expanded antigen-specific TCR sequences among all cell states. Each row corresponds to one expanded clone, clones that are shared between molecular states will exhibit a positive fraction in multiple columns. **(G)** Scatter plot showing the lack of a potentially confounding correlation between the fraction of CD8 T cells in the TEMRA state, and the sample collection time since onset. Each dot represents one donor and is colored by disease state.

