## Supplementary Methods for "Multimodal characterization of antigen-specific CD8^+^ T cells across SARS-CoV-2 vaccination and infection"

#### Human participants and PBMC collection

PBMC were collected from observational studies of adults (Supplementary Table 1) who were receiving BNT162b2 vaccination and willing to participate, excluding individuals with severe anemia or inability to comply with procedures. All groups were provided with written consent for enrollment with approval from the New York University (NYU) Institutional Review Board (across protocols 18-02035, 18-02037, and 12-01137). Participants had blood drawn at 4 defined time points, as diagrammed in Figure 1A, with 1-2 days flexibility in scheduling. Sample size calculations were not performed before the start of these nonrandomized, non-interventional studies, and outlier analyses were not performed.

Venous blood was collected by standard phlebotomy (total volumes ranging 40 - 80 mL). Within 5 hours of room temperature transport from an outpatient clinic, PBMCs were isolated from heparin vacutainers (BD Biosciences), followed by processing using SepMate (STEM-CELL Technologies), Ficoll-Paque Premium with density 1.077 (Cytiva), and Hank's balanced salt solution (ATCC), in accordance with manufacturers' recommendations. Aliquots of 1 mL were slowly frozen overnight within Corning CoolCell containers placed in -80 °C freezers, with cells suspended in complete media (RPMI 1640 supplemented with 40% fetal bovine serum) along with 10% DMSO, and after 2 days all vials were transferred to liquid nitrogen.

#### Flow cytometry and sorting

For initial CITE-seq and ASAP-seq experiments, PBMCs from all timepoints (Day 0, 2, 10, and 28) across 3 donors (12 specimens in total) were simultaneously thawed and promptly transferred to a 96-well V-bottom plate. This enabled further processing in parallel with multichannel pipettes. The same workflow was repeated with 3 additional donors, to generate the aggregate data in Figures 1-2. Each aliquot of 1-3 million frozen PBMC was thawed into 10 mL complete media, centrifuged at 300 RCF for 10 minutes at 4 °C, and resuspended in 200 μL conventional cytometry buffer (PBS with 4% fetal bovine serum), DAPI, and 2mM EDTA. Samples were passed through a 70-micron filter, and single cells were sorted on a FACSAriaII (BD Biosciences) using a 100-micron nozzle. The instrument operated via FACSDiva software, with post-sort analysis performed on FlowJo 10.8.1 (Tree Star). Gating excluded cellular debris and doublets based on FSC and SSC profiles and excluded dead cells based on DAPI. Cells were collected into 5mL of complete media separately maintained on ice until all sorting concluded, at which point all tubes were simultaneously centrifuged. Individual pellets were resuspended with 100 μL of staining buffer (PBS with 2% BSA and 0.01% Tween) along with unique hashing antibodies, followed by incubation on ice for 15 minutes. Hashed samples were washed 3 times with 500 μL of staining buffer and then pooled together. Viability (greater than 92%) and final cell counts were assessed with trypan blue and Countess II FL automated counter (ThermoFisher).

**CITE-seq library preparation**

Workflows for CITE-seq and cell hashing were performed as previously described[25,63]. An aliquot of 300,000 sorted and hashed cells was stained with 173 TotalSeq-A antibody panel (Supplementary Table 2). After incubating on ice for 30 minutes, cells were washed 3 times with 1 mL staining buffer to remove excess antibody. Cells were passed through a 40-micron Flowmi filter, resuspended in PBS, and ultimately loaded onto four lanes of 10x Genomics Chip G, following manufacturer protocols.

RNA library construction was performed according to the 10x scRNA-seq protocol, while the ADT and HTO library constructions were conducted following the CITE-seq protocol (https://citeseq.files.wordpress.com/2019/02/cite-seq_and_hashing_protocol_190213.pdf). During cDNA amplification (Step 2.2a), 0.2 µM of ADT additive primer (5′CCTTGGCACCCGAGAATTCC) and 0.1 µM of HTO additive primer (5′GTGACTGGAGTTCAGACGTGTGCTC) were added to the reaction mixture to enrich antibody tags. During cDNA cleanup (Step 2.3), supernatant containing the antibody tags was saved and further purified with 2x SPRI. The eluate was split into two tubes for ADT and HTO libraries. After cDNA cleanup, additional PCR reactions generated ADT and cell hashing libraries. These reactions were set up with KAPA Hifi Master Mix with the following primers: 10 µM 10x Genomics SI-PCR primer (5′AATGATACGGCGACCACCGAGATCTACACTCTTTCCCTACACGACGCTC), and 10 µM Illumina TruSeq DNA D7xx primer (5′CAAGCAGAAGACGGCATACGAGATxxxxxxxxGTGACTGGAGTTCAGACGTGTGC) for HTO library. 10 µM 10x Genomics SI-PCR primer, and 10 µM TruSeq Small RNA RPIx primer (5′CAAGCAGAAGACGGCATACGAGxxxxxxxxGTGACTGGAGTTCCTTGGCACCCGAGAATTCCA) for ADT library. The PCR products were purified with 1.6x SPRI.

**scATAC-seq library preparation**

ASAP-seq was conducted as previously described[26], with minor modifications. After staining with cell surface antibodies, cells were fixed in 0.1% formaldehyde for 5 minutes at room temperature. After washing, the cell pellet was resuspended in 100 µL of lysis buffer (10 mM Tris-HCl pH 7.4, 10 mM NaCl, 3 mM $MgCl_2$, 0.1% Tween-20, 0.1% Nonidet-P40 substitute (IGEPAL), 1% BSA) and kept on ice for 5 mins. The permeabilized cells were then resuspended with 1× Diluted Nuclei Buffer (10x Genomics) to a concentration of around 5000 cells/µL. 10 µL transposition mix (3µl 10x ATAC Buffer B and 7µl 10x ATAC Enzyme) was mixed with 5 µL sample and incubated for one hour at 37 °C. 0.5 µM bridge oligo A (TCGTCGGCAGCGTCAGATGTGTATAAGAGACAGNNNNNNNNNVTTTTTTTTTTTTTTTTTTTTTTTTTTTTT/3InvdT/) was added to the barcoding mix for proper amplification of antibody tags. The GEM incubation was performed with the following PCR program: 40 °C for 5 min, 72 °C for 5 min, 98 °C for 30 s; 12 cycles of 98 °C for 10 s, 59 °C for 30 s and 72 °C for

1 min; ending with hold at 15 °C. The post GEM incubation cleanup and library construction were conducted following the ASAP-seq protocol (https://citeseq.files.wordpress.com/2020/09/asap_protocol_20200908.pdf).

**Dextramer validation with spectral flow cytometry**

We initially tested a panel of 16 commercially available dextramer reagents (Immudex, catalog: RX19) designed to bind SARS-CoV-2 spike protein MHC class I epitopes[43] across 7 HLA haplotypes. All reagents were tagged with a unique DNA oligo barcode as well as PE fluorochrome. PBMC aliquots from all 4 timepoints for each donor were thawed as above and were subsequently resuspended in a cytometry buffer containing 0.1 g/L of herring sperm DNA (ThermoFisher) and Human TruStain FcX block (Biolegend). Cells were maintained in this blocking solution for 10 minutes at room temperature, 1 μL of each test dextramer reagent was subsequently added to each time-point sample, wells were thoroughly mixed, and the plate was incubated at 4 °C in the dark for 10 minutes. A separate antibody staining panel was also prepared in cytometry buffer, containing CD8a at 1:250 dilution, as well as 1:100 dilutions of CD2, CD4, CD14, CD16, and CD20. This was directly added (100 µl/well) to each well after initial dextramer incubation, wells were mixed, and the plate was returned to darkened 4 °C for 30 minutes. The plate underwent 4 rounds of centrifuge at 300 RCF 4 °C followed by wash with cytometry buffer, with final resuspension including DAPI and EDTA, followed by 70-micron filter passage. Samples were analyzed on a Cytek Aurora cytometer (Cytek) via SpectroFlow software (v3.03), with careful pre-calibration of fluorochrome spectral profiles to maximize accuracy and sensitivity. The gating strategy included: FSC, SSC; DAPI-negative; singlets; Dump-negative (CD14, CD16, CD20); CD2-positive; CD4-negative; CD8-positive; and a final dextramer/PE-positive gate to identify antigen-specific cells. Consistent with previous reports[53], only a subset of the 16 dextramer reagents exhibited an acceptable minimal non-specific binding at Day 0 and Day 2 time points, along with distinctly increased binding at Day 28 time point for the same test donor (see Supplemental Figure 5A).

We chose to employ 5 dextramer reagents that met this validation criteria, spanning the HLA-A*0201 and HLA-B*0702 alleles. These were loaded with the following spike (S) glycoprotein-derived immunodominant peptides, and tagged with respective DNA barcodes: VLNDILSRL with TTGTACTGAGTAAGC; YLQPRTFLL with CGGTTACAGTCGGTG; RLNEVAKNL with TCCAGGAACCATATG; NLNESLIDL with CGGTGTTAACGCGTT; SPRRARSVA with AGCTACTCGCACCAC. Our experiments also included a negative control reagent harboring the HLA-A*0201 allele loaded with a nonsense peptide (with barcode CAACTAATATGGTTA), as well as a nonsense HLA loaded with a nonsense peptide (with barcode GCAGACTTAGAAGAA). We identified 8 donors who stood out in exhibiting sizable antigen-specific T cell populations exclusively from Day 28 specimens (binding one or more of the 5 validated experimental dextramers), and utilized these samples to enrich for spike-specific $CD8^+$ T cells.

**Enrichment of spike-specific CD8$^+$ T cells, prior to ECCITE-seq**

To facilitate the study of rare populations, we enriched for spike-specific CD8$^+$ T cells prior to performing ECCITE-seq analyses. We aimed to facilitate enrichment while also mitigating the effect of potential biases, including the fact that no dextramer panel can successfully identify all spike-specific cells across all possible clonotypes. We proceeded to sort 3 populations: all dextramer-bound CD8$^+$ T cells (Bin 1), all CD38$^+$CD8$^+$ T cells (Bin 2), and an unenriched sampling of all CD8$^+$ T cells (Bin 3). Given the relatively scarcity of dextramer-positive cells, we enriched for this population first, and then obtained cells from the subsequent bins.

We stained Day 28 specimens with an aggregate panel of all 5 dextramers and 2 negative control reagents. A PCR tube was first loaded with 1.4 µL of 100uM d-Biotin (Thermo Fisher) diluted in PBS (to minimize non-specific binding). Then 10 µL of each dextramer specificity was sequentially added, the panel was well mixed, and ultimately 8.93 µL of this dextramer panel was added to each well of PBMC (consistent with manufacturer's recommended concentrations). A similar antibody panel as above (CD14, CD16, CD20, CD2, CD4, CD8) was added after dextramer, now also including CD38 at 1:100 dilution, as well as individual CITE-seq antibodies targeting CD8 and CD38. Final incubation with dextramers, fluorochrome antibodies, CITE-seq antibodies, and hashing antibodies ensued for 30 minutes at 4 °C in the dark. Subsequent cell preparation followed our prior cytometry protocol, except samples were loaded onto FACSAriaII for sorting. Gating was the same as above, with an additional CD38-high population created off the CD8 parent gate.

Since dextramer-positive CD8$^+$ events were the rarest, we collected all possible cells from this gate. Subsequently, we collected cells from the second and third bins. We then mixed all three bins together, at approximately 10% (Bin 1), 65% (Bin 2), 25% (Bin 3) ratio. This mixed pool was used as input for ECCITE-seq.

**ECCITE-seq library preparation**

Sorted cells were centrifuged at 400 RCF for 8 min at 4 °C and then resuspended in staining buffer. TotalSeq-C human cocktail (BioLegend) (Supplementary Table 2) was added for the surface protein staining, on ice for 30 min. After washing three times with 1 ml staining buffer, cells were resuspended in PBS and the cell concentration was adjusted to about 2000 cells/µL. Cells were loaded onto the 10x Chromium Next GEM Chip N, following manufacturer recommendations (Chromium Next GEM Single Cell 5' HT Reagent Kits v2). During cDNA amplification, 0.2 µM each of ADT (5′CCTTGGCACCCGAGAATT∗C∗C) and HTO (5′GTGACTGGAGTTCAGACGTGTGC∗T∗C) were added to the reaction. RNA, HTO, ADT and TCR libraries were constructed following the same protocols as above.

### Sequencing

Sequencing libraries were pooled and sequenced on an Illumina Novaseq using sequencing read lengths of 107 bp (read 1), 8 bp (i7 IndexRead), 16 bp (i5 IndexRead), and 107 bp (read 2). bcl2fastq was used to demultiplex raw sequencing data.

### Pre-processing, quantification, and QC of sequencing data

Sequencing data from ADT and HTO libraries were both aligned and quantified with salmon alevin (v1.8.0)[64]. Custom ADT and HTO indices, based on the DNA oligo barcode sequences, were constructed by running "salmon index" command. Single-cell barcode quantification matrices were generated by running "salmon alevin" command with the following parameters: -- naiveEqclass, --keepCBFraction 1.0. RNA sequencing data was aligned to the GRCh38 human reference genome using Cell Ranger (v6.0.0, "cellranger count") with default settings. ATAC sequencing data was aligned to the GRCh38 human reference genome using Cell Ranger ATAC (v2.0.0 "cellranger-atac count") with default settings. TCR sequencing data was aligned to the GRCh38/Ensembl human reference using Cell Ranger (v6.0.0, "cellranger vdj") with default settings.

For QC, we retained cells that passed the following thresholds: For the RNA modality, we retained cells that surpassed 500 UMI, and exhibited < 15% of reads mapping to mitochondrial regions. For the ATAC modality, we retained cells exhibiting at least 900 unique fragments per cell. For the ADT and HTO modalities in CITE-seq, we retained cells that surpassed 500 and 40 unique counts per cell, respectively. For the ADT and HTO modalities in ASAP-seq, we retained cells that surpassed 100 and 40 unique counts per cell, respectively. For each experiment, we retained cells that passed the required thresholds for each measured modality (i.e., for CITE-seq data, we retained cells that surpassed thresholds for RNA, ADT, and HTO modalities). After performing QC, we identified and removed doublets based on the cell hashing libraries, using the HTODemux function in Seurat[28] with default parameters.

### Visualization and clustering of CITE-seq data

To perform clustering and annotation of the original CITE-seq dataset (Figure 1B), we first processed the RNA and ADT modalities separately, performing normalization, dimensional reduction, and data integration steps. Subsequently, we performed weighted-nearest neighbor analysis[28], to jointly define cellular state based on RNA and protein data.

*Normalization and dimensional reduction*

We first split the CITE-seq data into 24 separate groups based on the combination of donor identity (n=6) and experimental timepoint (n=4). We performed normalization, feature selection, and dimensional reduction on each group independently.

For the RNA modality, we performed normalization using sctransform v1[65], using the SCTransform function in Seurat. This procedure also performs variance stabilization. We performed dimensional reduction using principal components analysis (PCA), retaining 40 dimensions. For the ADT modality, we performed normalization using the centered log-ratio (CLR) transformation, implemented in Seurat using the NormalizeData function with the arguments: normalization.method='CLR', margin=2. We centered the values for each feature to have a mean of 0 across all cells, but did not scale features to have unit variance, using the ScaleData function in Seurat (arguments: center=TRUE, scale=FALSE). We included all 173 ADT features for downstream analysis. We performed dimensional reduction using principal components analysis (PCA), retaining 40 dimensions.

*Data integration across donors and timepoints*

We next applied our 'anchor-based' data integration workflow[27] to integrate datasets produced across donors and timepoints. We performed separate integration analyses on both the RNA and ADT modalities. For the RNA modality, we selected a consensus set of 3,000 variable features across the 24 experimental groups using the SelectIntegrationFeatures command in Seurat. We performed integration as previously described using the 'reciprocal PCA' workflow, as implemented using the FindIntegrationAnchors (arguments: dims=1:40, reduction='rpca') and IntegrateData (default parameters) functions. This procedure returns a single 40-dimensional space (integrated.rna) that groups together shared cell states across donors and timepoints based on their transcriptomes. For the ADT modality, as also performed integration using the reciprocal PCA workflow, using all features and utilizing 40 dimensions. This procedure returns a single 40-dimensional space (integrated.rna.pca) that groups together shared cell states across donors and timepoints based on their protein data.

*Data integration across modalities*

To define cell state based on a weighted combination of RNA and ADT modalities, we constructed a weighted nearest neighbor (WNN) graph[28]. We constructed the graph using the FindMultiModalNeighbors (arguments: reduction.list=c("integrated.rna.pca', "integrated.adt.pca"), dims.list=c(1:40,1:40)) function in Seurat. The output of this procedure represents a cell graph ("wsnn") that was used as input for UMAP visualization, and graph-based clustering. We performed UMAP visualization using the RunUMAP command in Seurat with default parameters, and clustering using the FindClusters function in Seurat (arguments to FindClusters: graph.name="wsnn", resolution = 1). We performed differential expression on all pairs of clusters for both RNA and protein markers, and merged clusters that did not exhibit clear evidence of separation, or where the only differentially expressed features represented ribosomal genes or mitochondrial genes. In some cases (particularly for extremely rare cell types that required a higher resolution to be correctly annotated in our clustering), we increased the granularity of our clustering by subsetting cells in an individual cluster, and rerunning FindClusters on this subgraph.

**Differential cell type abundance analysis using Milo**

To identify differentially abundant cell states between Day 0 and Day 28, we utilized Milo[30] to analyze a weighted nearest neighbor graph generated from CITE-seq data. The precomputed shared nearest neighbor graph ("wsnn") was first used as input required for Milo using the "buildFromAdjacency" function (k=20, d=30). Next, cells were assigned into representative neighborhoods by running the "makeNhoods" function (refined=TRUE, prop=0.1, refinement_scheme="graph"). Cells were counted in neighborhoods using "countCells" function. To test for differential abundance, the "testNhoods" function was run (fdr.weighting = "graph-overlap") with design = ~batch + timepoint. Neighborhoods with SpatialFDR < 0.1 were determined as statistically significant for differential abundance, and were colored in Figure 1D-E.

**Gene module score**

To examine the strength of interferon response, we downloaded the list of genes that up-regulated in response to alpha and gamma interferon proteins from GESA website (https://www.gsea-msigdb.org/). We used the 'AddModuleScore' function in Seurat to quantify the expression of this gene module in single cells. In Figure 1C, Supplemental Figure 1C and D, one donor was excluded due to aberrant interferon expression at Day 28.

In order to identify a module of genes that were biomarkers of vaccine-induced cells, we performed differential expression analysis. We used the "FindMarkers" command in Seurat to compare expression of levels of vaccine-induced group A cells with CD8_TEM_3 cells (the most similar $CD8^+$ T cell cluster at level-3 resolution). We selected the top 200 genes (ranked by adjusted p-value) with adjusted p-value < 0.001 and minimal logFC threshold > 0.2. To ensure that our module was not contaminated by cell-cycle genes, we conservatively removed three genes that exhibited minimal up-regulation in group_A cells, but were strongly up-regulated in group B cells. The resulting 197-gene list is included in Supplemental Table 3.

**Mapping of ASAP-seq data with bridge integration**

To analyze the ASAP-seq dataset (Figure 2), we utilized our recently developed 'bridge integration' workflow[36], which integrates datasets that measure different modalities (i.e., scATAC-seq and scRNA-seq data) based on a 'bridge' dataset, where both modalities are measured simultaneously (i.e. a 10x multiome dataset). We downloaded a publicly available multiome dataset from 10x Genomics (https://www.10xgenomics.com/resources/datasets/pbmc-from-a-healthy-donor-granulocytes-removed-through-cell-sorting-10-k-1-standard-2-0-0), consisting of 11,351 paired scRNA-seq and scATAC-seq profiles of human PBMC, and utilized this as a bridge dataset to annotate each of our 78,677 ASAP-seq profiles.

To perform annotation, we followed the steps detailed in the cross-modality reference mapping Seurat vignette (https://satijalab.org/seurat/articles/bridge_integration_vignette.html), utilizing our CITE-seq dataset (Figure 1B) as a reference, and our ASAP-seq dataset as a query. The output of the bridge integration procedure includes multi-level cell annotations for each ASAP-seq profile, and additionally, visualizes the ASAP-seq dataset alongside our previously CITE-seq derived UMAP embedding.

We also performed further downstream analysis of the ASAP-seq dataset, based on the cell annotations derived from bridge integration. For these analyses, we performed TF-IDF normalization using the RunTFIDF function in Signac[66] with default parameters. We used normalized values to calculate 'gene activity' scores, which serve as a proxy for expression levels based on the average chromatin accessibility within and upstream of a gene body, using the GeneActivity function in Signac. To identify differentially accessible peaks in vaccine-induced cells, we used the "FindMarkers" function in Seurat, utilizing a logistic-regression based test[67] (arguments, test.use = 'LR', latent.vars = 'peak_region_fragments'), including cell-specific fragment count information to alleviate differences in cellular sequencing depth. The full list of differential peaks is included in Supplemental Table 4. We also used the top 1,000 differential peaks from this group as input to the FindMotifs function in Signac, which identifies enriched motifs from the JASPAR2022 database in this peak set compared to a background control set with matched GC content.

**Analysis of influenza vaccine ATAC-seq data**

We downloaded and re-analyzed publicly available scATAC-seq data[37] of samples before and after vaccination with the trivalent inactivated seasonal influenza vaccine (TIV) from GEO (GSE165906). We performed the same pre-processing steps as performed on our ASAP-seq dataset, using the 10x Genomics cellranger-atac software to align to the hg38 genome. One sample (donor id: 79) was excluded as an outlier from downstream analysis due to a low unique fragment number per cell. We integrated the ATAC modality across biological samples from different donors and timepoints. We applied reciprocal LSI projection to find integration anchors by running the "FindIntegrationAnchors" function in Seurat (reduction = "rlsi", dims = 2:30). The final integration was conducted using the "IntegrateEmbeddings" function to integrate the LSI coordinates across the datasets, returning a single 30-dimensional space (integrated_lsi). The integrated_lsi dimenstion of 2 to 30 were used as input for graph-based clustering, cell annotation, and UMAP visualization. To compare pseudobulk profiles of cells before and after vaccination, we quantified genomic bins using the "GenomeBinMatrix" function in Signac (arguments: binsize = 5000), retaining bins with at least one count.

**Visualization, clustering, and annotation of ECCITE-seq data**

Each ECCITE-seq profile simultaneously measures RNA and ADT modalities, but also measures immune repertoire sequences (TCR), as well as quantitative levels of the five MHC I

Dextramers loaded with SARS-CoV-2 spike peptides. To analyze this dataset, we utilized WNN analysis to jointly define cell state based on three modalities: integrated RNA, integrated ADT, and TCR. We also independently classified each cell as $Dex^+$ or $Dex^-$. Cells were classified as $Dex^+$ if the UMI counts for any of the five spike protein dextramers were at least two times as high as the UMI counts for the negative control. We annotated each TCR clone as 'spike-specific' if any individual cell in the clone was annotated as $Dex^+$.

Performing WNN analysis on multiple modalities requires a reduced-dimensional space to be independently generated for each modality. For RNA and ADT modalities, we generated this graph using the same normalization, data integration across samples, and dimensional reduction steps as we performed in our CITE-seq WNN analysis. In order to learn a separate low-dimensional space based solely on TCR sequences, we used clonotype neighbor graph analysis (CoNGA[68]), which employs the TCRdist distance metric[69] in order to quantify the similarity between two cells based on shared TCR sequence features. The script "setup_10x_for_conga.py" was first run in CoNGA with "--no_kpca" flag to prepare input files. The script "merge_samples.py" was run next to merge the datasets from multiple 10x lanes. By running the "run_conga.py" script with default settings, we performed kernel principal components analysis (kPCA) based on the TCRdist distance matrix and retained 40 components for downstream analysis. We used the three dimensional reductions (integrated RNA, integrated ADT, TCR) to perform a trimodal WNN analysis, which returned a single neighbor graph that integrated data from all three modalities. This graph was used as input for UMAP visualization, clustering, and annotation (Figure 3B).

We also annotated individual T cells as belonging to rare, small, medium, large, or hyperexpanded clones using the scRepertoire[70] package. The clonotype was called using the combination of the amino acid sequence of the CDR3 region for both the TCRα and TCRβ chains. The available chain was used for cells where only one of the two chains could be identified. For cells with multiple expressed chains, only the top two expressed chains were included for downstream analysis. We assigned clonal size for each cell by running the "clonalHomeostasis" function in scRepertoire with the proportional cut points cutpoints: (Rare = 1e-04; Small = 0.001; Medium = 0.01; Large = 0.1; Hyperexpanded = 1).

We compared each TCR with publicly available databases of T cells specific for SARS-CoV-2 peptide. We pooled TCRβ sequences from the ImmuneCODE COVID-19 TCR database[45] and the VDJdb COVID-19 TCR database[46]. When comparing TCR from our vaccination dataset, we restricted our overlap analysis to spike protein epitopes.

**Analysis of publicly available SARS-CoV-2 vaccination and infection datasets**

We downloaded a public vaccine CITE-seq dataset[24] from GEO (GSE171964), and mapped these data using our previously described 'reference-based mapping' workflow[27]. Our CITE-seq dataset was used as the reference, and RNA data from the public CITE-seq was used as the query. After identifying the anchors by running the "FindTransferAnchors" function in Seurat, the query data was projected onto the reference UMAP with the transferred cell-type labels using "MapQuery" function.

We obtained publicly available scRNA-seq dataset of acute SARS-CoV-2 infection samples[53] from (https://zenodo.org/record/5770747). The UMAP in Figure 4A is a reproduction of the visualization in the original manuscript. For further analyses, we used data from two individual sample sets: (1) patients CoV2_T001- CoV2_T010, acute; (2) patients CoV2_T011-CoV2_T020, acute. We retained cells with at least 500 detected UMI, mitochondrial read percentages lower than 15%, and where SNP-based demultiplexing was consistent with a single donor. As in the original manuscript[53], we removed a particular dextramer (peptide QYIKWPWYI) in the downstream analysis due to high nonspecific binding. As in the original manuscript[53], cells were labeled as CoV2-Dex$^+$ when the UMI count of a CoV2-Dextramer was higher than 10 and the fold change versus the negative control was more than five.

We obtained publicly available datasets from the COvid-19 Multi-omics Blood ATlas (COMBAT) Consortium[57], profiling human PBMC samples across multiple human donors at different stages of infection using ECCITE-seq (https://zenodo.org/record/6120249). We considered CD8$^+$ T cells from healthy donors and patients with mild, severe or critical symptoms. Cells with fewer than 300 detected genes or mitochondrial read percentage higher than 10% were removed. Donors including less than 200 CD8 T cells after QC were excluded. To perform integration across samples and modalities, we ran the same anchor-based integration procedure separately on the RNA and ADT modalities as we ran for our CITE-seq dataset. The WNN graph was generated using 30 RNA and 20 protein dimensions. The WNN graph was used as input for UMAP visualization and clustering.
